# Leveraging human-trained neural networks for cross-species chromatin regulation annotations

**DOI:** 10.1101/2025.10.16.682871

**Authors:** Noémien Maillard, Julie Demars, Raphaël Mourad

**Affiliations:** GenPhySE, Université de Toulouse, INRAE, ENVT, F-31326, Castanet Tolosan, France; MIAT, INRAE, F-31326 Castanet-Tolosan, France; University of Toulouse, UPS, 31062 Toulouse, France

**Keywords:** chromatin accessibility, histone marks, transcription factors, livestock species

## Abstract

Analogous to the Encyclopedia of DNA Elements (ENCODE) project, the Functional Annotation of ANimal Genomes (FAANG) consortium has produced chromatin annotations for domesticated animals, albeit in smaller amounts. Although acquiring experimental data is more accessible and affordable for many species, human and mouse organisms will remain the reference. Classical methods based on sequence conservation can be used to infer missing annotations, but are inappropriate for non-conserved sequences. While regulatory sequences share low to moderate conservation, they have retained their regulatory function during the evolution process. Here, we take advantage of three neural networks (DeepBind, DeepSEA, and Enformer) trained with human and mouse ENCODE data to infer chromatin annotations (transcription factors binding, chromatin accessibility, and histone marks) in cattle, pig, chicken, and European seabass. For this purpose, we comprehensively assessed the quality of predictions using experimental data from FAANG, through AUC-ROC and AUC-PR metrics. Our results showed similar predictions for various annotations in mammals and chicken, with AUC-PR ranging from 0.157 to 0.765 for H3K4me1 and H3K4me3, respectively, but lower in fish. Further analyses focused on pigs highlighted (i) accurate predictions even for non-conserved sequences, and (ii) variable predictions depending on genomic feature annotations. Our results advocate the widespread use of human-trained neural networks as a first step in cross-species genome annotation before training species-specific models.

## Introduction

The chromatin, the complex between the DNA double helices and proteins, regulates the organization of DNA in three dimensions within the nucleus over time. Interactions between transcription factors (TF), insulators, histones, and their modifications, and DNA modulate chromatin condensation, leading to a continuum of DNA accessibility degrees that participate in gene expression regulation^1^. Therefore, the characterization of chromatin regulation is required to improve the annotation of genomes for a better understanding of noncoding genetic variation in complex traits^2^.

For this purpose, the Encyclopedia of DNA Elements (ENCODE) consortium was created in the 2000s in order to generate data and help to elucidate how this regulation is modulated in model organisms, including human^3^ and mouse^4^. To date, almost 8000 human and 1400 murine ChIP-seq of both TF and histone marks, DNase-seq, and ATAC-seq experiments are referenced in the ENCODE dataportal^5,6^ (Supplementary Data 1). Later, in the 2010s, the Functional Annotation of ANimal Genomes (FAANG) initiative was organised to collect and analyse similar chromatin regulation data for species of agronomic interest, such as chicken, cattle, and pigs^7^. Initial efforts were made to standardize both protocols and computational analyses across different species, rather than conducting a large number of experiments. Although the FAANG dataportal is still in expansion through additional international projects^8^, much less data is available for livestock species (https://data.faang.org/). To date, 2,288 pig ChIP-seq and ATAC-seq experiments have been referenced, including biological and technical replicates for a limited number of experimental conditions (e.g., experiment type, tissue, developmental stage) (Supplementary Data 1).

As well as continuing to conduct costly and time-consuming wet lab experiments across species, computational methods based on sequence conservation and pairwise or multiple whole-genome alignment have been used over the last two decades to transfer annotations from well-annotated to poorly-annotated genomes^9,10^. However, multiple works have shown that non-coding sequences, including regulatory elements, often show weak conservation signals, thereby limiting the use of conservation-based tools^11,12^.

As an alternative to conservation-based methods, deep learning methods have been trained with ENCODE data to predict annotations of chromatin regulation, and a growing number of models are now available^13^. For example, transposable elements^14^ and regulatory activity predictions^15,16^ can be detected from sequence information. On the one hand, models are becoming increasingly sophisticated to improve performance through precision and recall metrics, while on the other hand, they are often used for a similar purpose in the study of model organisms^17^. Although chromatin regulation is conserved across large evolutionary distances^12,18–20^, few studies have evaluated the use of such novel tools for cross-species functional annotation by focusing on enhancer predictions^21–24^. Despite the affordable acquisition of omics experimental data in a broad range of species, human and mouse remain the reference, reinforcing the usefulness of these organisms for cross-species studies, including livestock.

Here, we comprehensively assessed the ability of three artificial neural networks (DeepBind^25^, DeepSEA^26^, and Enformer^27^), trained using human and mouse chromatin regulation data (ENCODE project, https://www.encodeproject.org/), to predict functional annotations from the reference genomes of livestock species, including pig, cattle, chicken, and seabass at the whole genome scale. We first compared the cross-species abilities of the different neural networks to predict a wide range of experiments. We then evaluated the performance of neural networks trained on human and mouse data to predict annotations across the phylogenetic tree and their limitations. Finally, we investigated the influence of sequence conservation and genomic features on the performance of chromatin prediction annotations.

## Results

### From both human and mouse-trained neural networks to cross-species predictions

To investigate the relevance of human and mouse-trained neural networks for translational studies, especially to identify genome-to-phenome links in livestock, we used three artificial neural networks presenting different properties (Supplementary Data 2) and trained on various biological experiments related to chromatin regulation (DeepBind^25^, DeepSEA^26^, and Enformer^27^) (Fig. 1a and Supplementary Data 2). DeepBind^25^ and DeepSEA^26^ are based on Convolution layers, while Enformer also uses Transformer layers^27^. In addition, the size of the input sequence ranges from 200 bp for DeepBind^25^, 2 kb for DeepSEA^26^, and up to 196 kb for Enformer^27^ (Fig. 1a and Supplementary Data 2). Cross-species predictions of functional chromatin annotations, including transcription factor binding sites and histone marks obtained from ChIP-seq data, as well as chromatin accessibility obtained from ATAC-seq and DNase-seq data, were performed using the reference genomes of the pig (Sscrr11.1), cattle (ARS-UCS1.2), chicken (bGalGal1) and European seabass (dlabrax2021) (Fig. 1b). The performance of the predictions was evaluated using standard metrics (area under the receiver operating characteristic curve [AUC-ROC] and area under the precision-recall curve [AUC-PR]) by comparing the predicted data with the observed data from the FAANG data portal (https://data.faang.org/) (Fig. 1c). AUC-ROC is classically used for benchmarks, but does not account for class imbalance (i.e. most bins do not overlap a peak). Hence, AUC-PR has also been used to account for class imbalance.

**Fig. 1:**
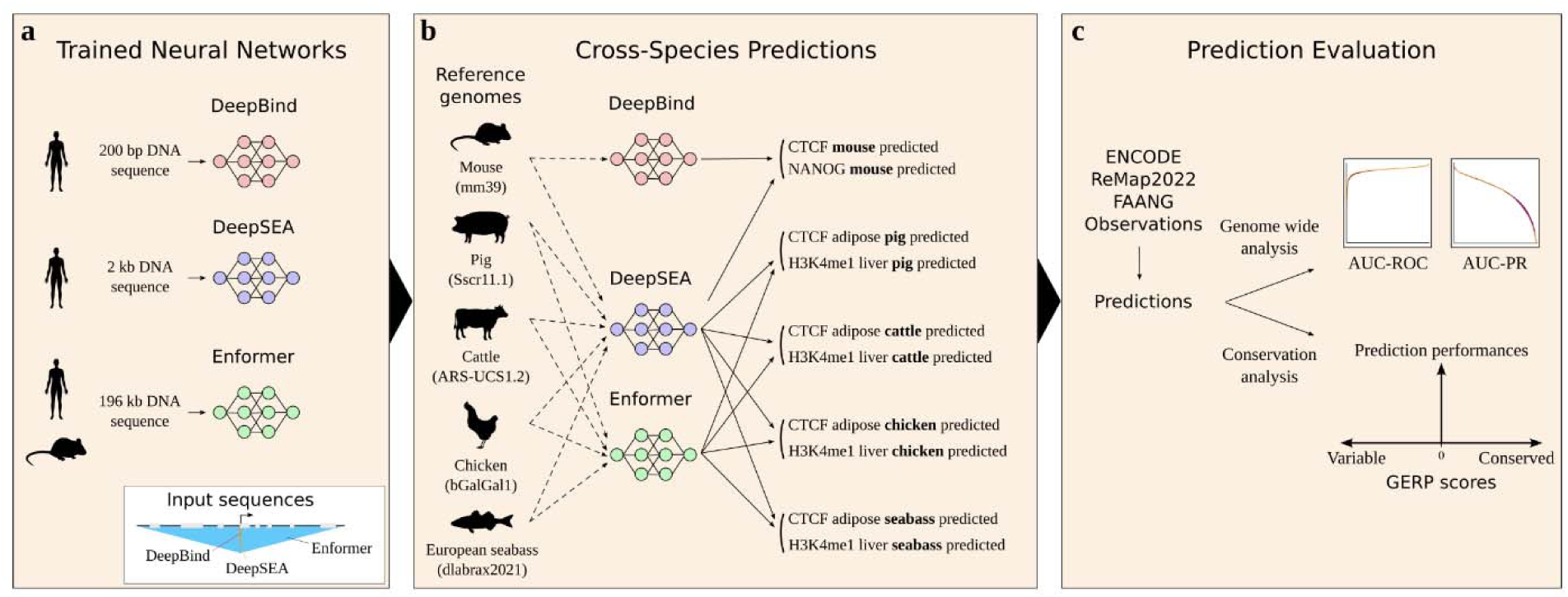
Approach to perform cross-species functional chromatin annotation predictions in livestock using human and mouse-trained neural networks. a. Neural networks trained on human and mouse data. Three neural networks are used, including DeepBind^25^, DeepSEA^26^, and Enformer^27^. b. Cross-species predictions. References genomes of mouse (mm39), pig (Sscr11.1), cattle (ARS-UCS1.2), chicken (bGalGal1), and European seabass (dlabrax2021) have been used. c. Evaluation of prediction performances. Standard metrics (AUC-ROC and AUC-PR) were calculated by comparing the predicted data with the observed data from the ENCODE (Encyclopedia of DNA Elements, https://www.encodeproject.org/), ReMap2022 (https://remap2022.univ-amu.fr/), and FAANG (Functional Annotation of Animal Genomes, https://data.faang.org/) databases. For a finer analysis, Genomic Evolutionary Rate Profiling (GERP) conservation scores have been used to show whether only sequences with conserved nucleotides can be well predicted.

### Prediction of chromatin annotations in mouse using human-trained models

To evaluate the ability of neural networks trained with human data to predict diverse annotations on the genome of another mammal, we first predicted annotations from the mouse reference genome (mm39) using only DeepBind^25^ and DeepSEA^26^. Since Enformer^27^ has been trained using both human and mouse data, predicting experiments from the mouse genome would likely lead to data leakage. We split the mouse genome into bins and then predicted the presence or absence of a ChIP-seq or DNase-seq peak in the bin. The comparisons included 9 transcription factors (TFs), 4 histone mark ChIP-seq, and DNase experiments (Fig. 2 and Supplementary Data 3). For human predictions, AUC-ROC and AUC-PR values were extracted from DeepBind^25^ and DeepSEA^26^ publications. For mouse data, 2 cell lines have been used as proxies according to the human training data (mouse B lymphocyte WEHI cell line, close to the human lymphoblastoid GM12801 cell line, used to train DeepBind CTCF model, and mESC equivalent to hESC used to train other DeepBind and DeepSEA models). For the binary classification, we considered a bin positive if at least half of the bin overlaps with an observed peak; otherwise, it was considered negative.

**Fig. 2:**
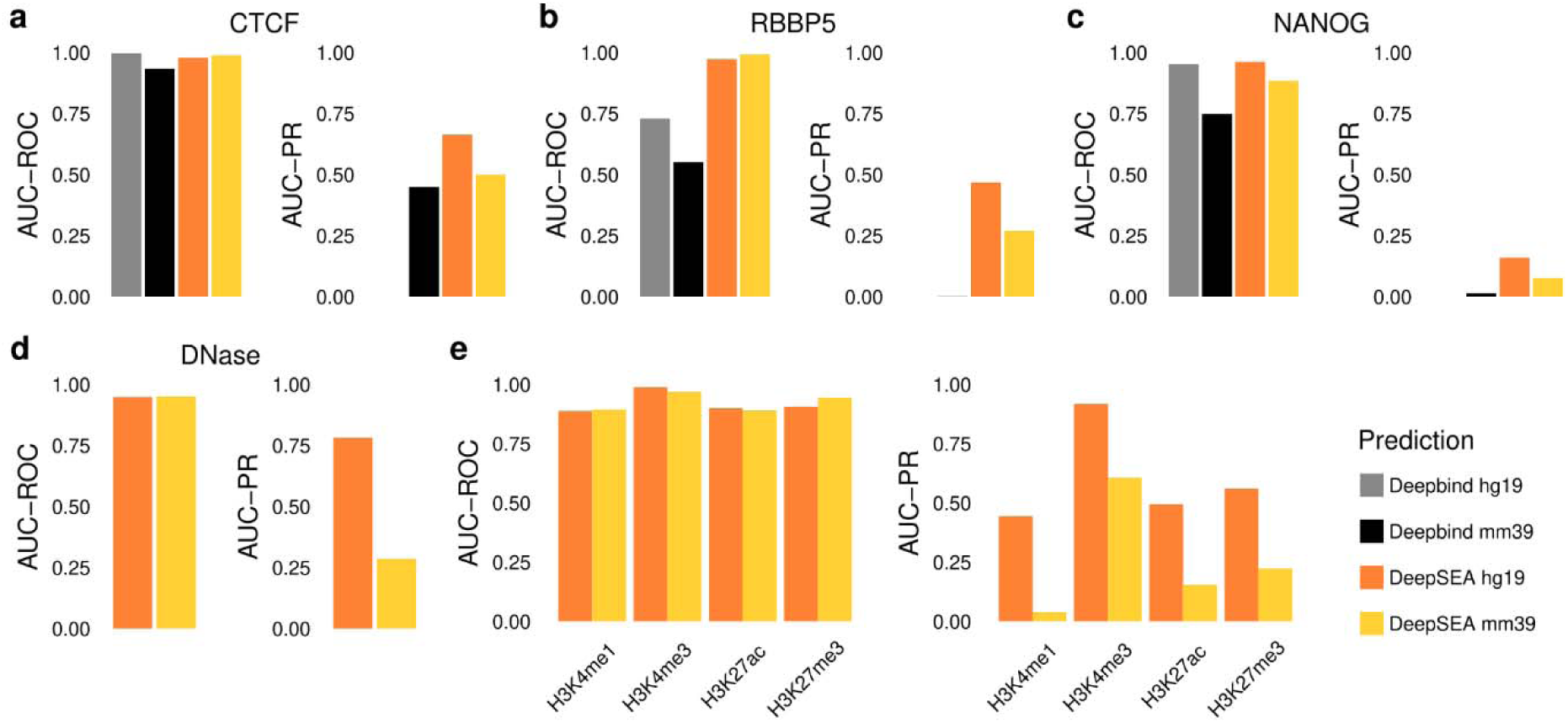
Predictive performances of two human-trained neural networks on the mouse genome. AUC-ROC (left) and AUC-PR (right) for CTCF (CCCTC-binding factor) (a), RBBP5 (RB Binding Protein 5, Histone Lysine Methyltransferase Complex Subunit) (b), NANOG (Nanog Homeobox) (c), DNase (d), and histone marks (e). The DeepBind prediction metric values are in grey and black for the human (hg19) and mouse (mm39) genomes, respectively. The DeepSEA prediction metric values are in orange and yellow for the human (hg19) and mouse (mm39) genomes, respectively. The human AUC values have been extracted from th original articles for DeepBind^25^ and DeepSEA^26^. Except for the DeepBind-CTCF prediction (model trained with GM12801 cell line), which was compared to CTCF conducted in WEHI cell line, the observations used for comparisons with predictions were conducted on mouse embryonic stem cells (ESCs) (n=1 for each comparison).

Predictions on the mouse genome showed high accuracy for CTCF peaks with AUC-ROC values of 0.934 and 0.989 using DeepBind and DeepSEA respectively, which were close to AUC-ROC values for the human genome (0.997 for DeepBind and 0.980 for DeepSEA) (Fig. 2a). For DeepSEA AUC-PR, which accounts for class imbalance, we obtained a value of 0.500 for mouse predictions, whereas it was 0.664 for human predictions (Fig. 2a). For all evaluated TFs, DeepSEA showed better AUC-PR values than DeepBind (Fig. 2b-c and Supplementary Data 3). For DNase and histone marks predicted only with DeepSEA, AUC-ROC values for mouse predictions were similar to those for human predictions, but there was a more significant gap between AUC-PR values between mouse and human predictions (Fig. 2d-e). In addition, we also observed that ubiquitous TFs, including CTCF (CCCTC-Binding Factor) and RBBP5 (RB Binding Protein 5, Histone Lysine Methyltransferase Complex Subunit), showed higher AUC-PR values than TFs involved in specific biological processes or developmental stages, such as NANOG (Nanog Homeobox) (Fig. 2a-c and Supplementary Data 3).

Overall, the correlations between human and mouse AUC-PR values appeared to be stronger for TF binding sites, regardless of the AUC-PR values, than for histone and DNase experiments, despite the greater amount of peak sequence in the training data (Fig. 3a). Indeed, when training a machine learning model, it is expected that the amount of training data (here the amount of peak sequences) will strongly influence prediction performances. In agreement, we observed a strong correlation between the number of peaks used to train the models and the AUC-PR calculated with DeepSEA predictions on both human (Fig. 3b) and mouse (Fig. 3c) genomes. We then focused on the *Sox2* and *Igf2* loci, which are known to carry binding sites for NANOG and CTCF, respectively (Fig. 3d). For NANOG predictions, 1 true positive out of 4 was predicted in the *Sox2* region. However, a high error rate has also been measured with 3 false negatives and 1 false positive (Fig. 3d). Conversely, CTCF experimental peaks are well predicted in the *Igf2* region, albeit at a high false positive rate (Fig. 3d). Like CTCF, DNase and H3K4me3 observed peaks were correctly predicted despite false positive peak predictions (Fig. 3d).

**Fig. 3:**
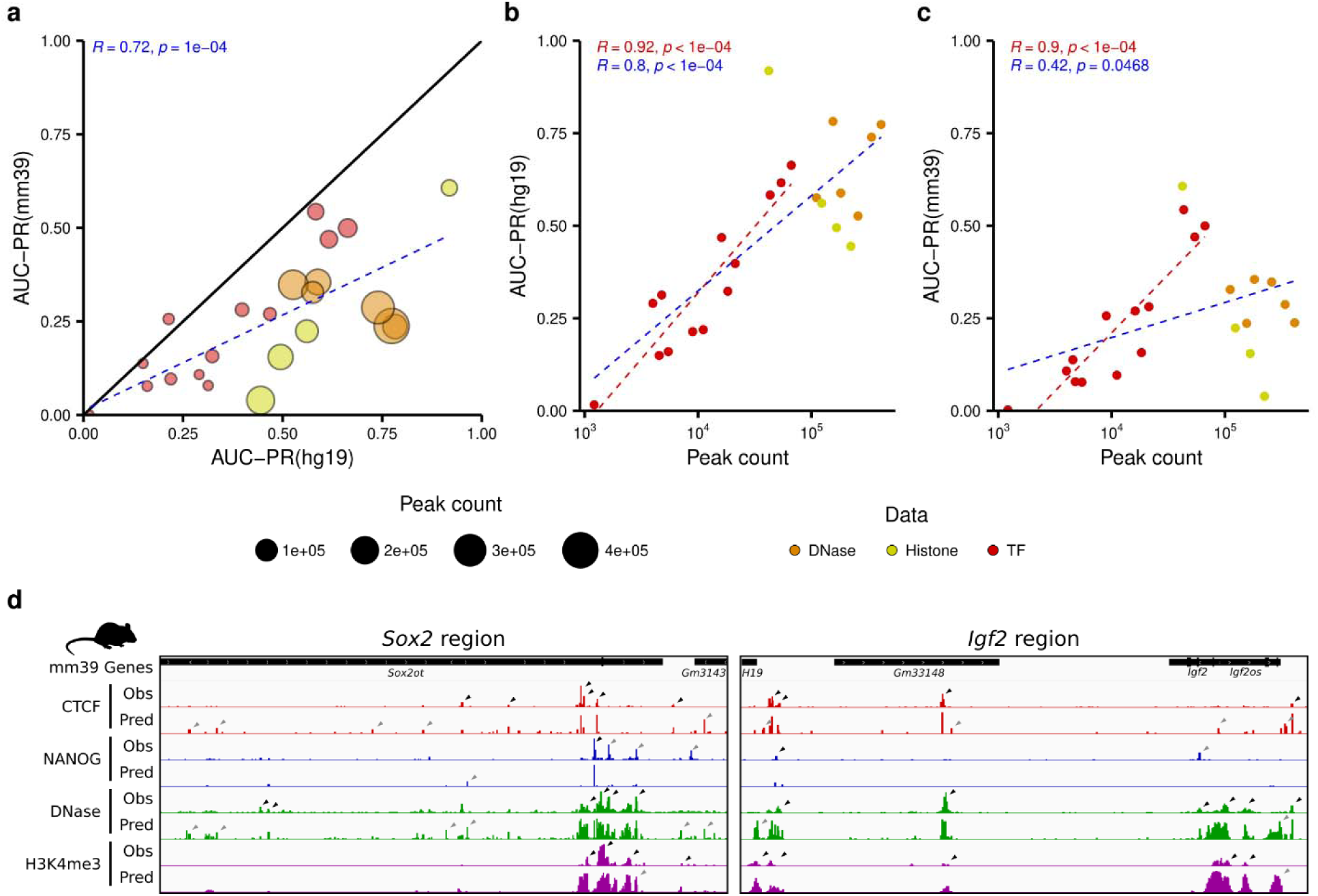
Comparison of the predictive performances of DeepSEA across different types of experiments. a. Correlation between human (hg19, x axis) and mouse (mm39, y axis) AUC-PR values (blue dotted line and statistics). The size of the dots indicates the number of peaks counted in the experiments used to train DeepSEA. b and c. Correlations between peak count of experiments used for DeepSEA training and AUC-PR of human (b) or mouse (c). The colors of the dots represent the type of experiments with transcription factor binding sites (red), DNase (orange), and histone marks (yellow). The correlations take only TF (red dotted lines and statistics) into account or all experiments (blue dotted lines and statistics). d. Visualization of the *Sox2* and *Igf2* regions through the Integrative Genome Viewer. The mouse annotated genes are shown on the first track with *Sox2* and *Igf2* on the left and right panels, respectively. For CTCF (red), NANOG (blue), DNase (green), and H3K4me3 (purple), both observed experiments in mouse embryonic stem cells (ESC) (top) and predicted annotations (bottom) are shown for comparison. Annotations of the mouse genome are from Ensembl release 112.

Taken together, our results demonstrated two levels of analysis depending on TF, or chromatin accessibility and histone mark predictions. For TF predictions, our results showed that the amount of data for training (i.e., the number of peaks) is a critical factor for accurate cross-species predictions, which makes it more difficult to predict TFs that are specific to a particular biological process. For chromatin accessibility and histone marks, the lower correlations with the amount of data sustain biologically more complex experiments.

### Prediction of chromatin annotations in livestock using human and mouse-trained models

To assess the predictive ability of the neural networks across the phylogenetic tree, we then predicted CTCF, four histone marks, and chromatin accessibility experiments in five tissues (adipose, liver, lung, muscle, and spleen) with DeepSEA^26^ and Enformer^27^ from reference genomes of livestock species. For the data analyses, the chosen predictions are mentioned in Supplementary Data 4. We chose two mammals (pig, Sscr11.1, and cattle, ARS-UCS1.2), one bird (chicken, bGalGal1), and one fish (European seabass, dlabrax2021). We used the DeepSEA model to compare the metrics between the human, the mouse, and the livestock species genomes. We also used the Enformer model, which is expected to outperform other neural networks given its additional use of Transformer layers (Fig. 4 and Supplementary Data 5). As for mouse prediction assessment, we considered a bin positive if at least half of the bin overlaps an observed peak; otherwise, it was considered negative.

**Fig. 4:**
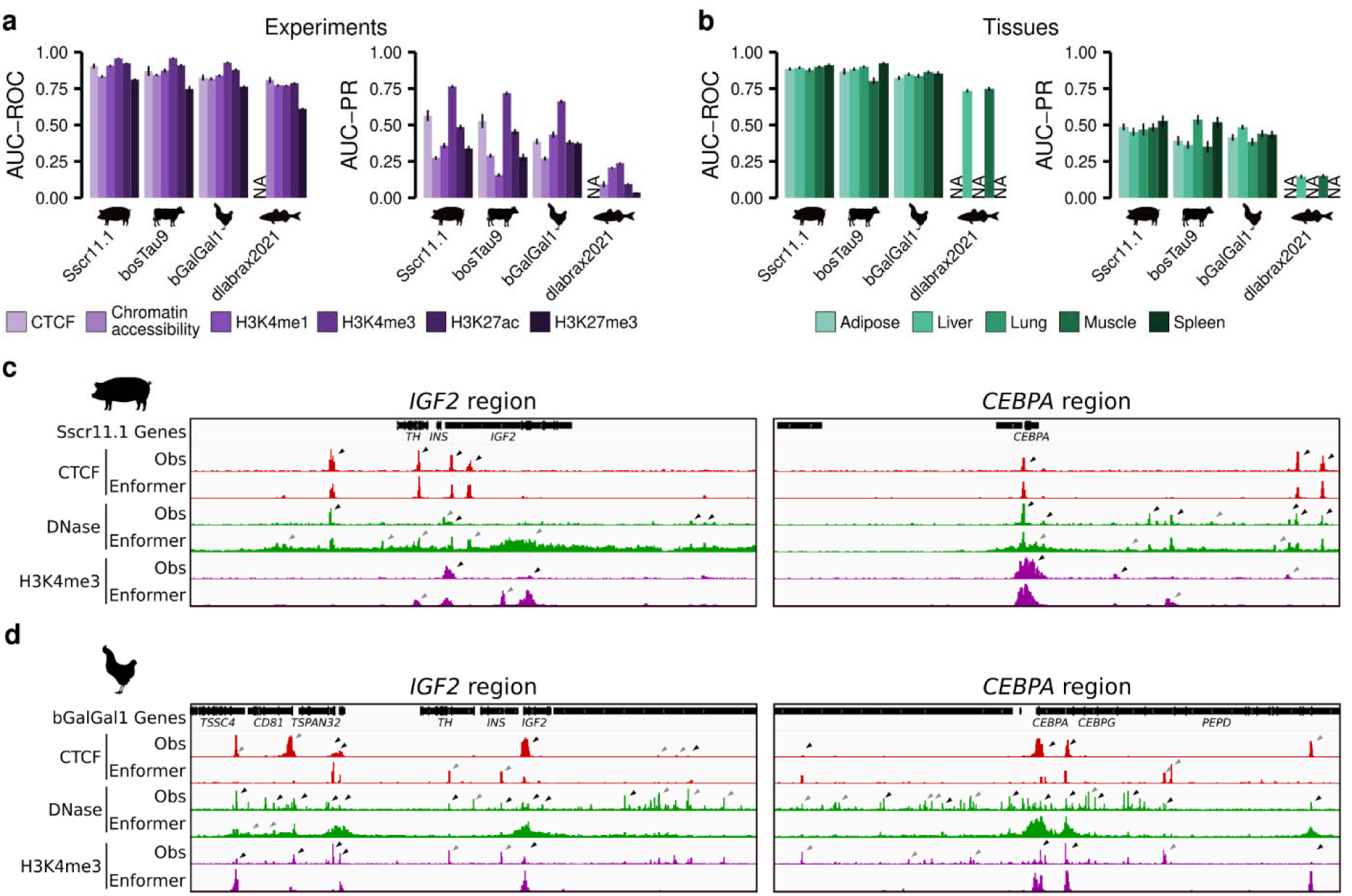
Predictive performances of the Enformer human and mouse-trained neural network on different livestock genomes. a and b. AUC-ROC (left) and AUC-PR (right) for the same predictions from pigs, cattle, chicken, and European seabass reference genomes with Enformer^27^ grouped by experiments (a, purple) or by tissue (b, green). Data shown are means ± SE. Grouped by experiments, n=36, 30, 44, 52, 34, and 44 for CTCF, Chromatin accessibility, H3K4me1, H3K4me3, H3K27ac, and H3K27me3, respectively. Grouped by tissues, n=50, 62, 34, 56, and 38 for adipose, liver, lung, muscle, and spleen, respectively. c and d. Visualization of the *IGF2* (left) and *CEBPA* (right) regions in pig (c) and chicken (d) through the Integrative Genome Viewer. The annotated genes are shown on the first track. For CTCF (red), DNase (green), and H3K4me3 (purple), both observed experiments in pig and chicken liver tissue (top), and predicted experiments (bottom) are shown for comparison. Annotations of pig and chicken genomes are from Ensembl release 112.

First, the AUC-PR metrics obtained using DeepSEA for the comparison between the mouse and the four species studied varied according to the experiments (Fig. 2a and 2d-e and Supplementary Data 5a). The binding of CTCF (0.500) and DNase (0.287) appeared to be better predicted in mice than in pigs (0.353 ± 0.035 se and 0.220 ± 0.012 se), cattle (0.309 ± 0.035 se and 0.230 ± 0.017 se), and chickens (0.284 ± 0.010 se and 0.230 ± 0.021 se). Similarly, H3K27me3 was better predicted in mice. These results can be the consequence of overall low predictive performance and the absence of replicates for the evaluation of predictions from the mouse genome. Conversely, H3K4me1, H3K4me3, and H3K27ac in pigs, cattle, and chickens seem to be predicted more accurately than in the mouse (Fig. 2d-e and Supplementary Data 5a). Second, Enformer showed better predictive ability than DeepSEA, since all AUC-PR values obtained with Enformer (Fig. 4a-b) are higher than those measured with DeepSEA (Supplementary Data 5a-b) for all four species and tissues.

Overall, we observed more variability in the prediction metrics across different experiments than between different tissues (Fig. 4a-b and Supplementary Data 5a-b). However, AUC-PR values are lower in cattle adipose, liver, and muscle tissues than in pig and chicken tissues, reflecting metabolic specificities of those tissues in ruminants (Fig. 4b). Indeed, monogastric and ruminant energetic metabolism evolved differently due to their respective energetic substrate (glucose and short fatty acids), as reviewed in^28,29^. Thus, our results seem to support this by showing particular chromatin regulation related to energetic metabolism in cattle^30^, which might be the consequence of no ruminant data in the training dataset. For non-mammalian genomes, although we found slightly less good metrics for the chicken genome, much lower metrics for the European seabass genome were found, suggesting a shift in the predictive ability of neural networks across phylogeny (Fig. 4a-b).

To illustrate predictions, we focused on two genomic regions: the *IGF2* locus, which undergoes genomic imprinting in pigs (Fig. 4c) but not in chickens (Fig. 4d)^31^, and the *CEBPA* region, which is a master transcription factor gene for adipogenesis^32^. Overall, as previously observed from genome-wide AUC-PR values, the Enformer neural network showed better predictive ability than the DeepSEA model (Fig. 4c-d and Supplementary Data 5c-d). For CTCF predictions in the IGF2 region, all peaks have been correctly predicted in pigs and half of the peaks in chickens (Fig. 4c-d). Conversely, more false positives are observed for H3K4me3 peaks in pigs than in chickens in both regions, while the opposite is shown for DNase (Fig. 4c-d).

Our results revealed that cross-species chromatin predictions based on neural networks trained on human data go beyond mammals, especially for CTCF, H3K4me3, and H3K27ac (Fig. 4 and Supplementary Data 5).

### Prediction performances regarding the phylogeny and sequence conservation

In order to better understand the relationship between predictive annotation metrics across species, we first evaluated the correlation between the AUC-PR of H3K4me3 and the phylogenetic distance calculated as substitutions per site (Fig.5a-c). The differences of DeepSEA AUC-PR values measured among the species seem explained by the phylogenetic distances (Fig. 5a), since strong correlations are obtained between the four livestock species, mouse and human (Fig. 5b). A similar trend is observed for the Enformer neural network (Fig. 5c). This suggests a breakpoint in prediction performance at a certain sequence evolution distance beyond 1.166 substitutions per site.

**Fig. 5:**
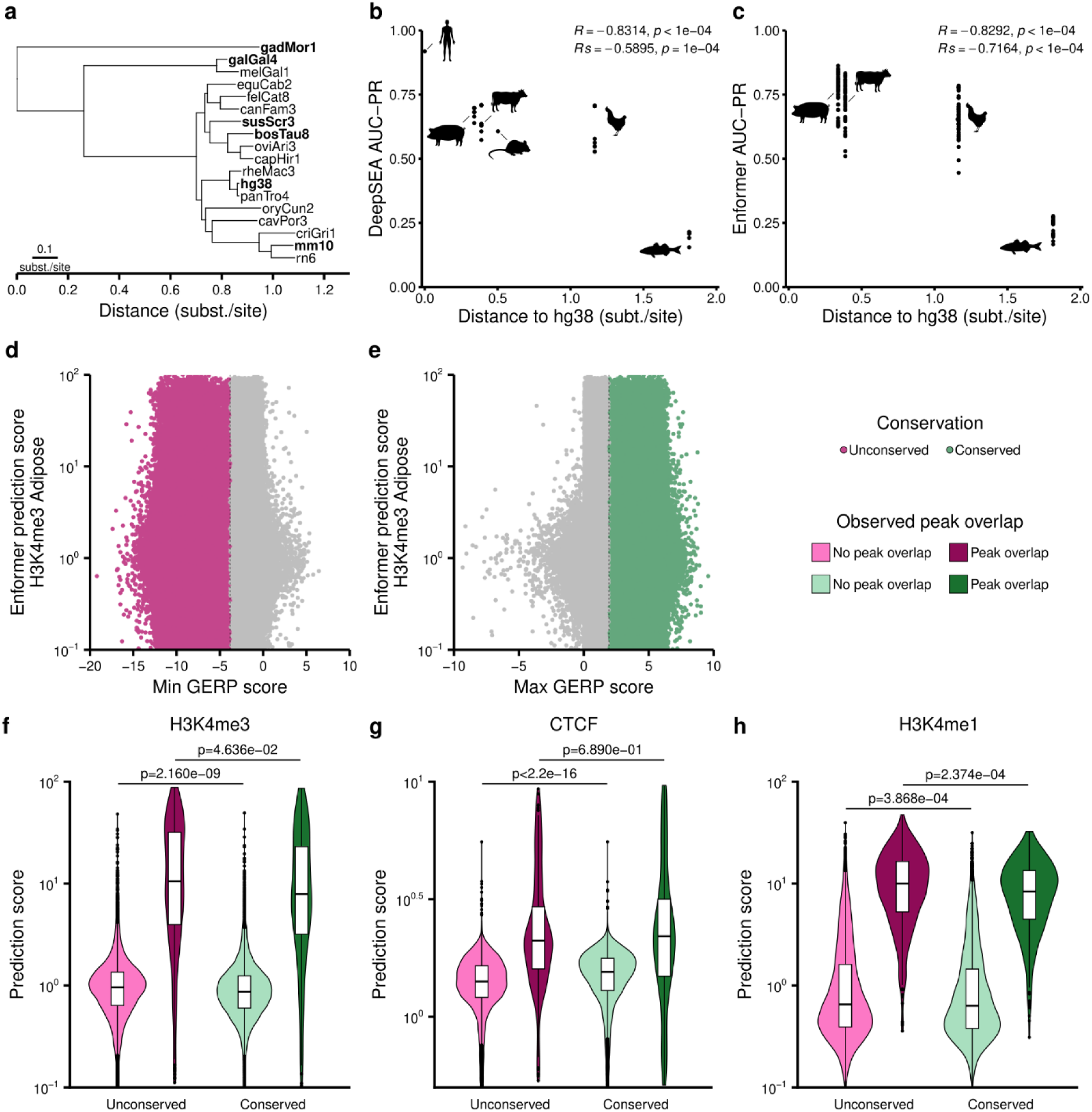
Impact of the sequence conservation on the performance of predictions in pigs. a. Phylogenetic distance (substitutions per site) of commonly studied species and species of interest (bold). gadMor1 is used as a proxy for European seabass. b and c. Relationship between AUC-PR of H3K4me3 predicted by DeepSEA^26^ (b) and Enformer^27^ (c) and phylogenetic distances, relative to the human genome, of species of interest. Pearson (R) and Spearman (Rs) correlations have been calculated, excluding human AUC-PR. d and e. Distribution of minimum GERP cores (d) and maximum GERP scores (e). 1% of bins have been sampled all along the pig reference genome (n=1,897,280). For each bin, the maximum and minimum GERP scores have been extracted. Vertical line is the mean of minimum (d) and maximum (e) GERP used to differentiate unconserved (pink, n=372,291) or conserved (green, n=372,267) bins. f to h. Distribution of Enformer prediction scores of H3K4me3 (f), CTCF (g), and H3K4me1 (h) in adipose tissue according to their conservation status (pink vs green) and the overlap (dark pink or green) or not (light pink or green) with an observed peak. For each experiment, a two-sided unpaired Wilcoxon test has been realized between unconserved not overlapping a peak bins (light pink, n=13781, 13960, 13565 for H3K4me3, CTCF and H3K4me1 respectively) and conserved not overlapping a peak bins (light green, n=13744, 13988, 13657 for H3K4me3, CTCF and H3K4me1 respectively), and between unconserved overlapping a peak bins (dark pink, n=394, 215, 610 for H3K4me3, CTCF and H3K4me1 respectively) and conserved overlapping a peak bins (dark green, n=407, 163, 494 for H3K4me3, CTCF and H3K4me1 respectively). Additionally, r effect sizes have been calculated for each group: r=-0.036, -0.070, 0.154, -0.021, -0,022 and -0.111 for H3K4me3 no peak overlap, H3K4me3 peak overlap, CTCF no peak overlap, CTCF peak overlap, H3K4me1 no peak overlap, and H3K4me1 peak overlap groups, respectively. A negative r effect size indicates a prediction score higher in the unconserved group, and a positive r effect size indicates a prediction score higher in the conserved group.

In addition, we used a metric for sequence conservation - the Genomic Evolutionary Rate Profiling (GERP) score - to investigate the impact of conserved and unconserved sequences on prediction performances (Fig. 5d-f). For H3K4me3, we down-sampled 1% of bins along the pig genome, and the maximum and minimum GERP scores were used per bin. We evaluated the distribution of these GERP scores in function of the prediction scores of each bin (Fig. 5d-e). We showed that a high proportion of nucleotides displaying negative GERP scores, reflecting low sequence conservation across species, presented high prediction scores as well as highly conserved nucleotides with positive GERP scores (Fig. 5d-e). To focus on the prediction scores, we extracted bins with a minimum GERP score lower than the mean of the minimum GERP scores (the least conserved). We did the same for bins with a maximum GERP score greater than the mean of maximum GERP scores (the most conserved). We then split the bins into two classes, depending on whether or not they overlapped with an observed peak (Fig. 5f). For H3K4me3, we show that a statistical difference exists between conserved and unconserved sequences, whether bins overlap an observed peak or not (p-value=0.0463 and 2.160x10^-9^, respectively, on a two-sided unpaired Wilcoxon test). However, the differences measured between classes were negligible (0.070 and 0.036 for bins overlapping a peak or not, respectively). CTCF bins overlapping a peak showed no statistical difference between conserved and unconserved groups, but this may be due to smaller sample sizes (Fig. 5g). In contrast, all other comparisons showed the same tendency as H3K4me3 (Fig.5f-h).

Overall, our results showed that cross-species neural network prediction accuracy goes beyond sequence conservation.

### Prediction performances in pigs with respect to genomic organization

We then assessed whether prediction performances depend on genomic organization. For this purpose, we predicted 6 experiments in 5 tissues on the pig reference genome and evaluated AUC-ROC and AUC-PR metrics depending on different genomic features (Fig. 6a).

**Fig. 6:**
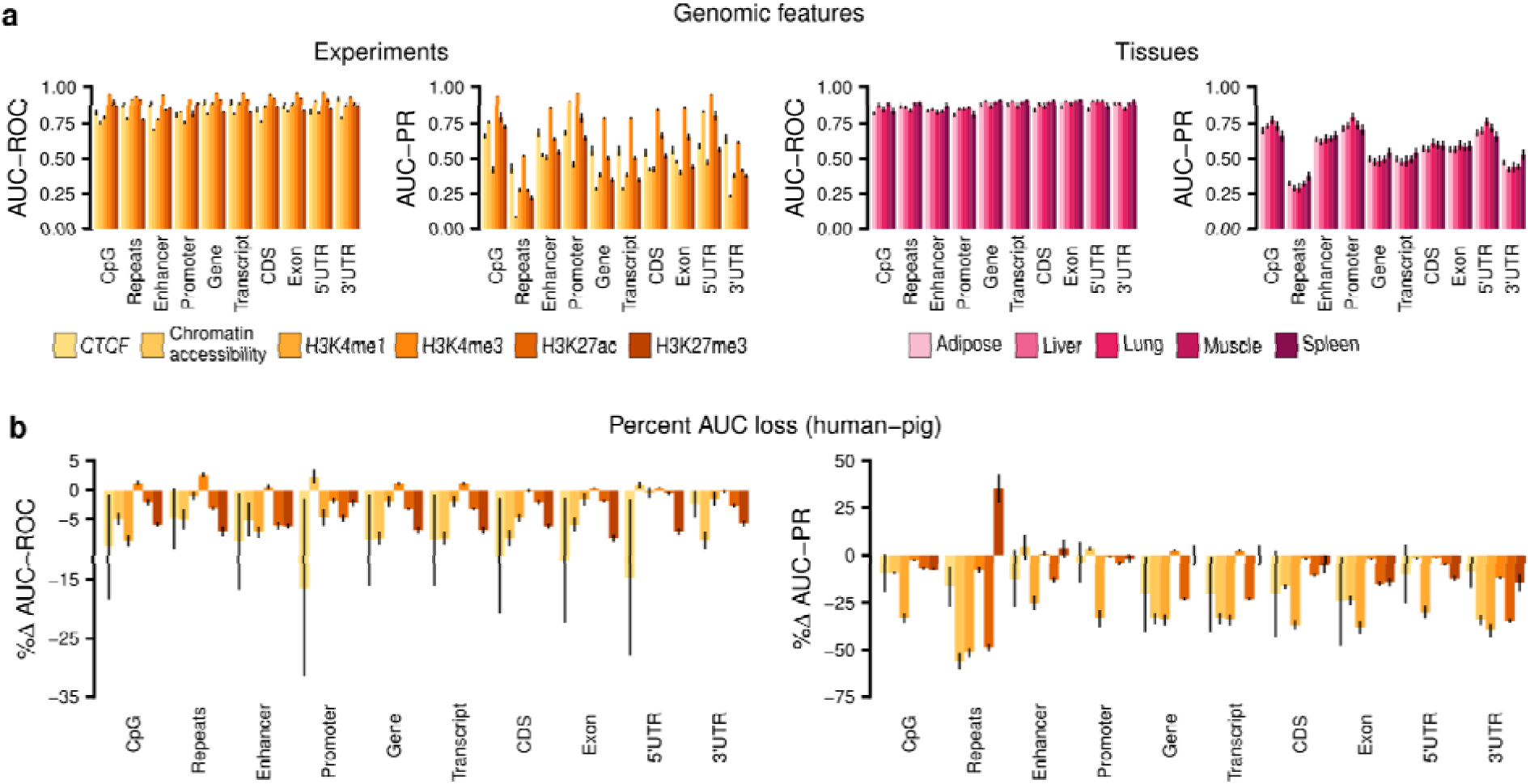
Impact of pig annotated genomic features on the performance of Enformer predictions. a. AUC-ROC (left) and AUC-PR (right) calculated for 8 genomic features between observations from the FAANG project and predictions from the pig reference genome with Enformer^27^, grouped by experiments (yellow) or tissues (pink). Data shown are means ± SE. Grouped by experiments, for each genomic feature, n=36, 30, 44, 52, 34, and 44 for CTCF, Chromatin accessibility, H3K4me1, H3K4me3, H3K27ac, and H3K27me3, respectively. Grouped by tissues, for each genomic feature, n=50, 62, 34, 56, and 38 for adipose, liver, lung, muscle, and spleen, respectively. b. Difference in percentage of AUC-ROC (left) and AUC-PR (right) from human (GRCh38/hg38) to pig (Sscr11.1) muscle. For each genomic feature, 1 human AUC has been subtracted from pig AUCs (n=2, 6, 12, 14, 10, and 12 for CTCF, Chromatin accessibility, H3K4me1, H3K4me3, H3K27ac, and H3K27me3, respectively). Data shown are means ± SE.

We found that promoters, CpG islands (CGIs), and 5’UTR regions are particularly well predicted with AUC-PR values above 0.564 for all experiments, except H3K4me1, which has a mean AUC-PR of 0.459 ± 0.158 SE, 0.422 ± 0.172 SE, and 0.474 ± 0.141 SE for promoters, CpG, and 5’UTR, respectively (Fig. 6a). In contrast, repeated sequences and 3’UTR regions showed AUC-PR under 0.521 for all experiments, except CTCF and H3K4me3 in 3’UTR, which cap respectively at 0.640 ± 0.221 SE and 0.616 ± 0.091 SE (Fig. 6a). With promoters, enhancers are the main chromatin regulatory features, but they display lower sequence conservation compared to promoters. Our results show that they are slightly less well predicted than promoters, CpG, and 5’UTR, but better predicted than the similarly predicted CDS and exons. Indeed, the results showed similar AUC-PR for H3K4me3, H3K27ac, and H3K27me3 marks for enhancers, exons, and CDS, but they showed better AUC-PR for CTCF (0.685 ± 0.192 SE, 0.562 ± 0.190 SE, and 0.543 ± 0.174 SE for enhancers, exons, and CDS, respectively), chromatin accessibility (0.526 ± 0.073 SE, 0.479 ± 0.076 SE, and 0.425 ± 0.071 SE), and H3K4me1 (0.510 ± 0.141 SE, 0.401 ± 0.136 SE, and 0.425 ± 0.143 SE) (Fig. 6a).

In addition, we measured the difference in prediction performance between human and pig genomes in muscle tissue (Fig. 6b). The results showed that the strongly conserved regulatory features (promoters, CpG, 5’UTR) seem to have the least AUC-PR loss. Moreover, the regulation pattern of promoter activity seemed well predicted by the neural networks, as shown by poor AUC-PR loss of H3K4me3 and DNase-seq. In contrast, the enhancers seem well predicted, but their pattern seems not fully understood (Fig. 6a). Indeed, the enhancer annotations from Ensembl are based on H3K4me1 and H3K27ac co-localization, but with AUC-PR rising to 0.510 ± 0.141 and 0.640 ± 0.090, respectively, our results show moderate predictions on those marks in the enhancers. While H3K27ac can be found not only in enhancers but also in promoters, H3K4me1 is mostly known as an enhancer-repressive mark. However, our results show that H3K4me1 is best predicted in enhancers over all features (Fig. 6a) and, with a delta AUC-PR of -25.0%, they also show the least loss from human to pig (Fig. 6b), suggesting a better association of this mark with enhancers. Those results are in concordance with DeepGCF^33^ functional conservation scores and suggest that neural networks improve the overall predictions.

Altogether, our results showed that prediction performances vary across genomic features, with promoters, CpG islands, and 5’UTR regions being the most accurately predicted, but results are encouraging for the analysis of less conserved sequences as enhancers.

## Discussion

Here, we demonstrated the accuracy of neural networks trained with human and mouse data to predict annotations on genomes of livestock species. Extensive benchmarking revealed great prediction performances on genomes from the same phylogenetic clade as the genome used for the training of neural networks (Mammals), but more strikingly, also for a more distant clade (Aves). However, the results suggested a breakpoint at a phylogenetic distance beyond 1.166 substitutions per site, since for the seabass genome, prediction metrics were very low.

A few studies have highlighted conservation of regulatory elements in mammals^19,34^, vertebrates^18,20^, and other animals^12^. These fundamental results demonstrate that the function of gene regulatory elements, such as enhancers, is maintained over the course of evolution regardless of their sequence conservation^35,36^. In parallel, during the last two decades, computational methods based on sequence conservation have been used for cross-species prediction, but are inherently not appropriate for unconserved sequences^10,37,38^. For example, GERP++ relies on the alignment of the genome sequences of many divergent species and searches for sequences whose similarity has been maintained during evolution. In contrast, more recently, machine learning methods, which do not rely on sequence conservation but instead on sequence features such as DNA motifs, have shown that enhancers share functional properties across species, supporting biological data, but this was often limited to a few species of mammals^22,23,33,39^. For instance, the two-step neural network approach, DeepGCF, first predicts epigenomic functional effects and then uses these effects as input to predict the functional conservation between humans and pigs^33^. Here, our findings are in concordance with the literature, supporting that we can predict chromatin regulation beyond mammals, regardless of sequence conservation.

Although we were able to evaluate the ability of neural networks to predict across several species and point out strengths as well as some limits, our analysis was limited by the data available in FAANG for the evaluated livestock species. It would be of great interest to evaluate such performances for an even broader range of experiments. For example, the evaluation of other histone modifications would have been interesting, such as H3K9me3, which has been shown to be enriched for silencing LTRs, LINEs, and SVAs repeated sequences, and with tissue specificities^40^. In addition, it would be valuable to study intermediate species across the tree of life, such as reptiles and amphibians, to accurately define the shift in chromatin prediction annotation abilities. However, accessing valuable chromatin-omics data for experimental purposes may present an obstacle to performing AUC-PR.

As pointed out here and by the literature, machine and deep learning models are not perfect, and several options exist in order to improve the cross-species predictions. Over Enformer and its use of Transformer layers, it has been shown that model architecture is a key element in highlighting different features and interactions between them^22,41^. Moreover, Kelley showed that using multi-species data during the deep learning model training phase improves predictions^42^, and while annotation data from not commonly studied species are often lacking, alternatives like pseudo-labeling have been proposed^43^. Finally, the improvement of molecular techniques used to study chromatin regulation and annotate genomes might have some biases (sequence affinity of enzymes, for example). Novel innovative methods, such as CUT&Tag, for efficient epigenomic profiling, that have not been used in the training set nor in the evaluation set, may likely improve predictions of chromatin accessibility^44^. Here, we used three artificial neural networks relying on different parameters (DeepBind^25^, DeepSEA^26^, and Enformer^27^) as a first step to evaluate their abilities for cross-species predictions of chromatin annotation, mainly in livestock species. This area of research is in constant evolution, and novel models are regularly released. As an example, ChromBPNet seems to show prediction performances close to Enformer performances but with a lighter architecture based on convolution layers^45^. While it has not been used for cross-species predictions, this suggests that other models might be used in order to improve cross-species predictions.

In summary, our research opens the field of predicting experiments for species of agronomical interest with artificial neural networks trained with human and mouse data. While neural networks are generally developed to predict variant impact in human pathologies, we show here that we can also use them for cross-species predictions and consider their use in the study of complex traits for species of agronomical interest.

## Methods

### DeepBind model

DeepBind^25^ is a convolutional neural network taking 200 bp sequences as input (http://tools.genes.toronto.edu/deepbind/). The output is a pseudo-probability of a given protein, ranging from 0 to 1. The network has been used to predict the binding of CTCF, RBBP5, NANOG, MAX, MYC, POU5F1, SIN3A, SUZ12, YY1 with the corresponding models, respectively D00328.018, D00797.001, D00786.001, D00504.005, D00785.001, D00795.001, D00804.002, D00812.001, D00710.007. DeepBind predictions have been computed on the Genobioinfo cluster of the Genotoul-Bioinfo platform, parallelized on 64 CPU cores (approximately 11 hours for the predictions with all 9 models on the whole mouse genome).

### DeepSEA model

DeepSEA^26^ is a convolutional-based model taking a 2000 bp (context and sequence) as input, and which outputs a pseudo-probability to observe a peak in a 200 bp centered sequence. The model has been downloaded from Kipoi. DeepSEA was used to predict 2002 experiments (DNase, ChIP-seq of transcription factors, and histone marks) for each sequence. Transcription factors (CTCF, RBBP5, NANOG, MAX, MYC, POU5F1, SIN3A, SUZ12, YY1), DNase, and histone ChIP-seq experiments (H3K4me1, H3K4me3, H3K27ac, H3K27me3) from tissues of interest (liver (HepG2), lung (A549), muscle (HSMM)) have been selected. DeepSEA predictions have been computed with an NVIDIA RTX4090 GPU (prediction times ranging from 50 minutes for the seabass genome to 2h30 for the pig and cattle genomes).

### Enformer model

Enformer^27^ is a deep learning model composed of convolutional layers followed by transformer layers. It takes a 196,608 bp length sequence as input and outputs 896 bins of 128 bp length. Enformer predicts 5313 human genomic tracks (DNase-seq, ChIP-seq of transcription factors and histone marks, ATAC-seq, and CAGE) for each sequence. We extracted ChIP-seq experiments of CTCF, H3K4me1, H3K4me3, H3K27ac, H3K27me3, DNase-seq, and ATAC-seq related to adipose, liver, lung, muscle, and spleen tissues. Enformer predicts a normalized read count. Enformer predictions have been computed with an NVIDIA RTX4090 GPU (prediction times ranging from 30 minutes for the seabass genome to 2h50 for the human genome).

### Mouse genome

Mouse reference genome (GRCm39, mm39) from Ensembl release 112 has been used to predict peaks with DeepBind and DeepSEA. For this purpose, the genome has been binned into 200 bp and 2000 bp bins for DeepBind and DeepSEA, respectively, and then DNA sequences have been extracted according to bins. For each sequence, the models were used to predict the pseudo-probability of binding for the different experiments.

### Mouse ChIP-seq and DNase-seq data

All mouse experimental data have been obtained in the embryonic stem cell line (ESC). ChIP-seq bed files of transcription factors (CTCF (GSE60917 and GSE29218), RBBP5 (GSE22934), NANOG (GSE44286), MAX (GSE48175), MYC (GSE90893), POU5F1 (GSE44286), SIN3A (GSE81081), SUZ12 (GSE127804), YY1 (GSE68195)) have been extracted from ReMap2022^46^ and bed files of DNase-seq and histone mark experiments have been extracted from ENCODE (DNase-seq (ENCFF178MGP), ChIP-seq H3K4me1 (ENCFF426IIV), ChIP-seq H3K4me3 (ENCFF247GVM), ChIP-seq H3K27ac (ENCFF360VIS), ChIP-seq H3K27me3 (ENCFF008XKX)) (Supplementary Data 6). The peaks have been lifted over the more recent genome mm39 with the liftOver^9^ tool from UCSC (Supplementary Data 7).

### AUC-ROC and AUC-PR values extraction from publications

For human (GRCh37, hg19 for both DeepBind and DeepSEA) ChIP-seq peak predictions, we extracted available AUC-ROC and AUC-PR values from publications.

### AUC-ROC and AUC-PR computation

AUC-ROC and AUC-PR have been computed using the PRROC R package (version 1.3.1). For class labeling, a bin has been considered positive (class 1) if at least half of the bin overlaps with an observed peak; otherwise, it has been considered negative. For this, the findOverlaps function from the GenomicRanges package was used (version 1.56.2) with default parameters (maxgap=-1L, ignore.strand=TRUE) except minoverlap set at half of the bin width (100L for DeepBind and DeepSEA and 64L for Enformer).

### Livestock species genomes

Pig reference genome (Sscrofa11.1), chicken reference genome (GRCg7b, bGalGal1), and European seabass reference genome (dlabrax2021) have been downloaded from Ensembl release 112. The cattle reference genome (ARS-UCD1.2, bosTau9) has been downloaded from Ensembl release 99. Each livestock species’ genome has been used to predict peaks with DeepSEA and Enformer. Each genome has been binned into 2000 bp and 196,608 bp bins for DeepSEA and Enformer predictions, respectively. For each sequence, DeepSEA was used to predict the pseudo-probability to observe a peak, and Enformer predicted a normalized read count. For DeepSEA and Enformer predictions, AUC-ROC and AUC-PR have been calculated with the same method as for mouse experiments.

### Livestock species ChIP-seq and chromatin accessibility data

For each experiment, pig, chicken, and cattle data bed files have been downloaded from the FAANG consortium (GSE158430). ChIP-seq (CTCF, H3K4me1, H3K4me3, H3K27ac, H3K27me3), DNase-seq (chicken), and ATAC-seq (pig and cattle) performed in adipose, liver, lung, muscle, and spleen tissues have been chosen. European seabass bigBed of the same experiments in lung and spleen have been downloaded from the AQUA-FAANG consortium (ensembl.org database), then bed files have been created from bigBed using bigBedToBed from UCSC (Supplementary Data 6). The bed files have been kept as is, except for the chicken data that were lifted over to the more recent genome bGalGal1 using the liftOver^9^ tool from UCSC.

### Pig genome annotations

The GTF files of genomic feature annotations (Gene, Transcript, CDS, Exon, 5’UTR, 3’UTR) and regulatory annotations (promoter, enhancer) have been extracted from Ensembl database release 115. Repeats have been extracted from the UCSC database. CpG islands have been extracted from the UCSC Table Browser. For conserved regions analysis, GERP scores have been extracted from Ensembl release 113 (Supplementary Data 7).

### Human genome

To calculate AUC-ROC and AUC-PR differences from human to pig with Enformer predictions, the Human reference genome (GRCh38, hg38) from Ensembl release 112 has been used. For this purpose, the genome has been binned into 196,608 bp bins for Enformer, and then DNA sequences have been extracted according to bins. For each sequence, the model was used to predict the normalized read count for the different experiments. AUC-ROC and AUC-PR have been calculated with the same method as for mouse experiments.

### Human genome annotations

The GTF file of genomic feature annotations (Gene, Transcript, CDS, Exon, 5’UTR, 3’UTR) and regulatory annotations (promoter, enhancer) have been extracted from Ensembl database release 115. Repeats have been extracted from the UCSC database. CpG islands have been extracted from the UCSC Table Browser (Supplementary Data 7).

### NGS read alignments

For Integrative Genome Viewer^47^ (IGV) panels, mouse, pig, and chicken raw reads have been aligned on mm39, Sscr11.1, and bGalGal1 reference genomes, respectively. Mouse raw reads have been extracted with the SRA run selector for transcription factors and from the ENCODE consortium for H3K4me3 and DNaseI-Hypersensitivity experiments. Pig and chicken raw reads have been extracted from data referenced in the FAANG consortium corresponding to GSE158430 (PRJEB14330, PRJNA665197, PRJNA665199, PRJNA665209, PRJNA665214, PRJNA665216, PRJNA665216). For ChIP-seq, inputs have been merged for each tissue-sample couple (Supplementary Data 8). For the alignment of raw reads, the nf-core ChIP-seq (for CTCF and H3K4me3 experiments) and ATAC-seq (for ATAC-seq and DNase-seq experiments) pipelines have been used with default parameters. The macs_fdr parameter has been set to 0.01 for ChIP-seq experiments and 0.05 for ATAC-seq/DNase-seq experiments.

### Integrative Genome Viewer

Integrative Genome Viewer^47^ (IGV) has been used to show how predictions fit observations for genomic regions. Predicted peak heights have been set with the autoscale parameter.

## Statistical information

Wilcoxon tests have been computed using a two-tailed *wilcox_test* function from the coin R library. The r effect size has been computed by dividing the Z statistic from the Wilcoxon tests by the square root of the sample sizes for each test.

## Code and data availability

All the available data from ENCODE, ReMap2022, and FAANG used in this article are mentioned in Supplementary Data 6-8.

All the code is available at: https://forge.inrae.fr/genesis/EAGLE.

All the bigwig prediction files are available at: https://entrepot.recherche.data.gouv.fr/ under the accession numbers https://doi.org/10.57745/TN3R6F, https://doi.org/10.57745/92QIQS, https://doi.org/10.57745/EEQDSP.

## Acknowledgements

We would like to thank Sarah Djebali, Laurent Bréhélin, Bertrand Servin, and Juliette Riquet for their help and advice. We would also like to thank Guillaume Devailly for his help in the R analysis.

## Funding

This study is supported by the PEPR Agro-écologie et Numérique PDOC2 – ANR-22-PEAE-0015 PEAE 2022 and by the DIGIT-BIO Metaprogram Research from INRAE.

## Contributions

N.M. performed analyses, wrote the manuscript, and drew figures. J.D. and R.M. conceived the study, secured funding, supervised N.M., and read and edited the manuscript.

## Competing interests

The authors declare no competing interests.

## Notes

https://forge.inrae.fr/genesis/EAGLE

https://doi.org/10.57745/TN3R6F

https://doi.org/10.57745/92QIQS

https://doi.org/10.57745/EEQDSP

### Competing Interest Statement

The authors have declared no competing interest.

### Summary of Updates

- figure 6 that have been updated with results from promoters and enhancers - additional supplementary data 4 to explain which data have been used for the training and the predictions - some editing mistakes

