## Supplementary data 1 to 3 and 5 for "Leveraging human-trained neural networks for cross-species chromatin regulation annotations"

### Slide 1
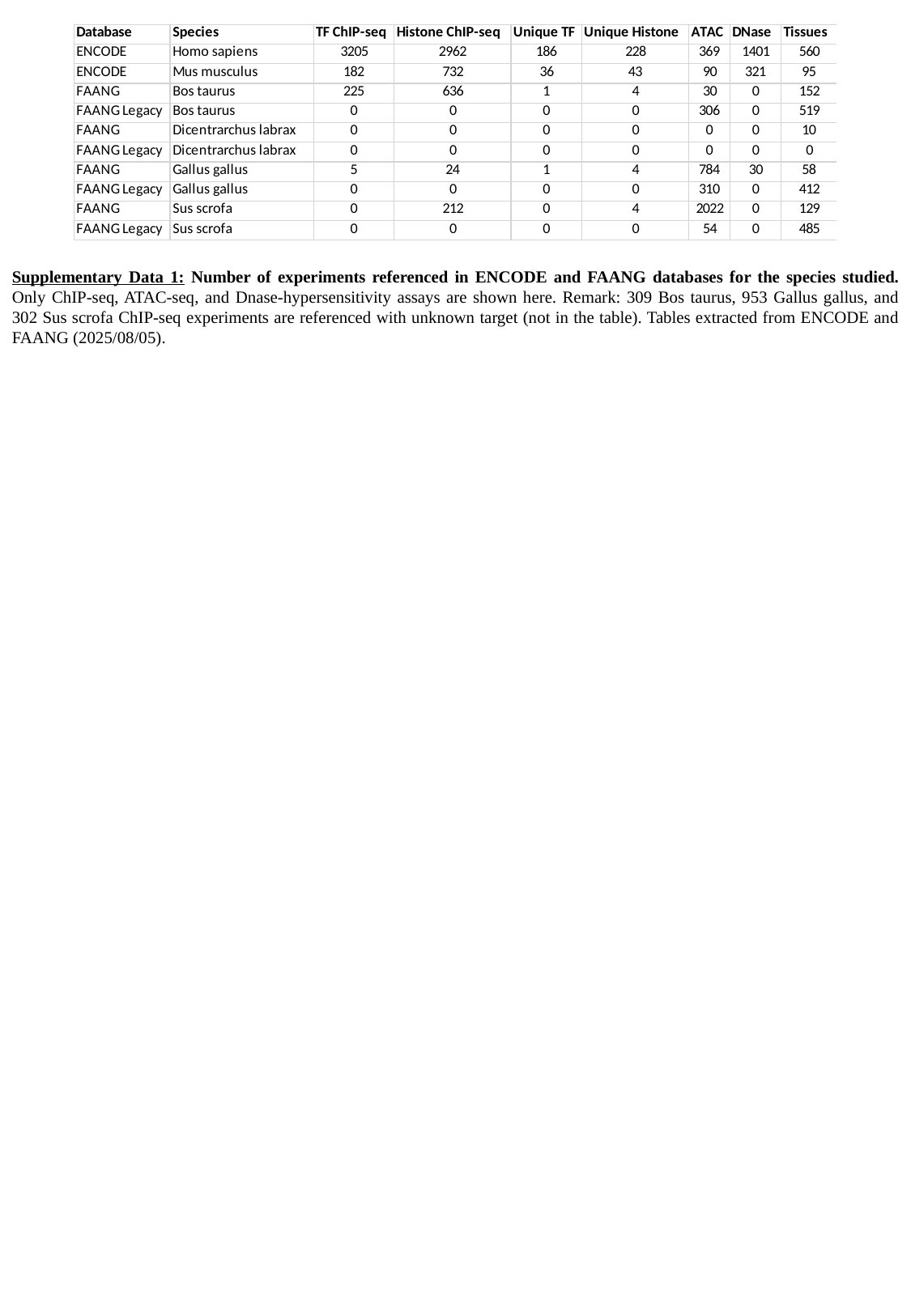

Supplementary Data 1: Number of experiments referenced in ENCODE and FAANG databases for the species studied. Only ChIP-seq, ATAC-seq, and Dnase-hypersensitivity assays are shown here. Remark: 309 Bos taurus, 953 Gallus gallus, and 302 Sus scrofa ChIP-seq experiments are referenced with unknown target (not in the table). Tables extracted from ENCODE and FAANG (2025/08/05).

### Slide 2
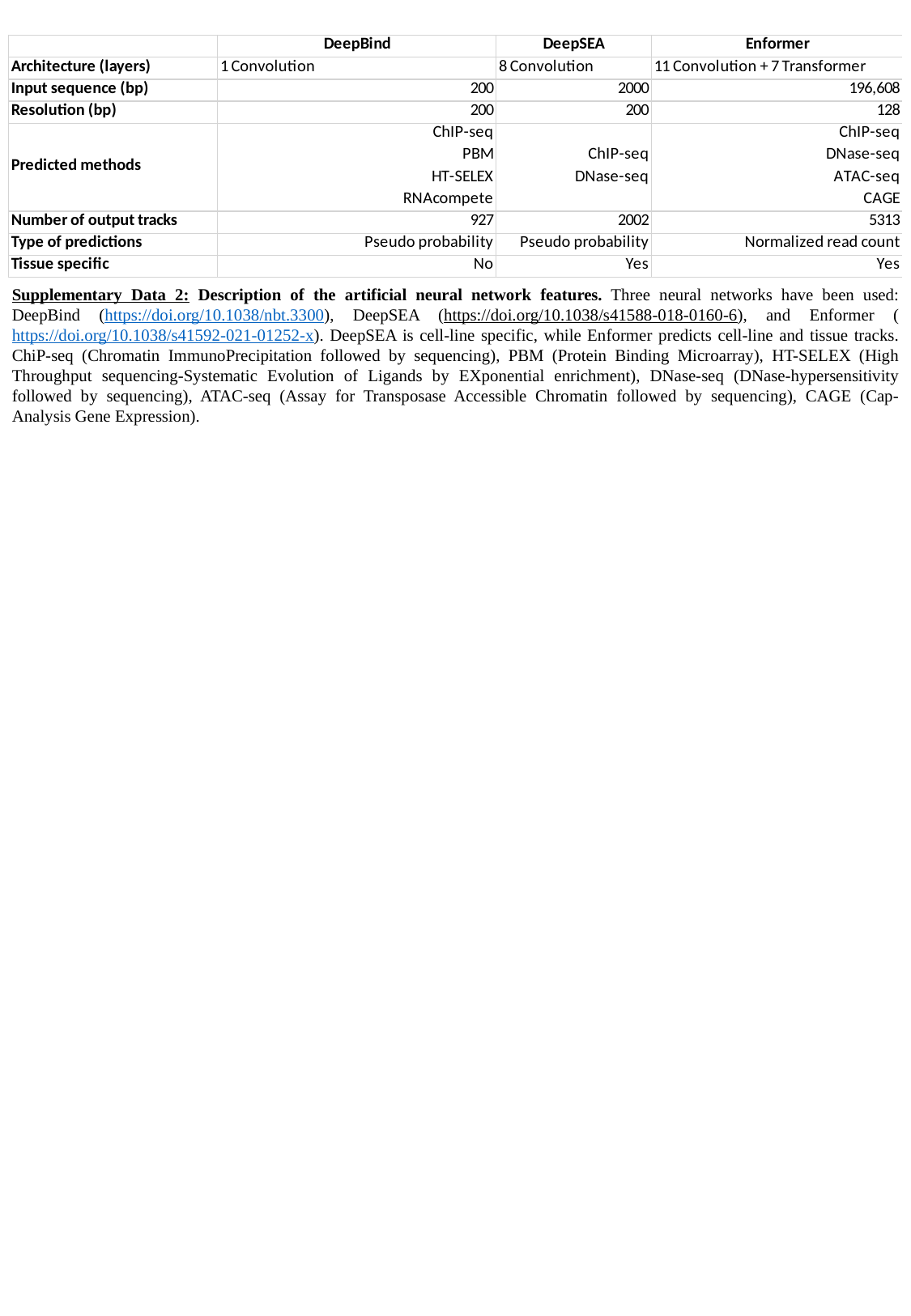

Supplementary Data 2: Description of the artificial neural network features. Three neural networks have been used: DeepBind (https://doi.org/10.1038/nbt.3300), DeepSEA (https://doi.org/10.1038/s41588-018-0160-6), and Enformer (https://doi.org/10.1038/s41592-021-01252-x). DeepSEA is cell-line specific, while Enformer predicts cell-line and tissue tracks. ChiP-seq (Chromatin ImmunoPrecipitation followed by sequencing), PBM (Protein Binding Microarray), HT-SELEX (High Throughput sequencing-Systematic Evolution of Ligands by EXponential enrichment), DNase-seq (DNase-hypersensitivity followed by sequencing), ATAC-seq (Assay for Transposase Accessible Chromatin followed by sequencing), CAGE (Cap-Analysis Gene Expression).

### Slide 3
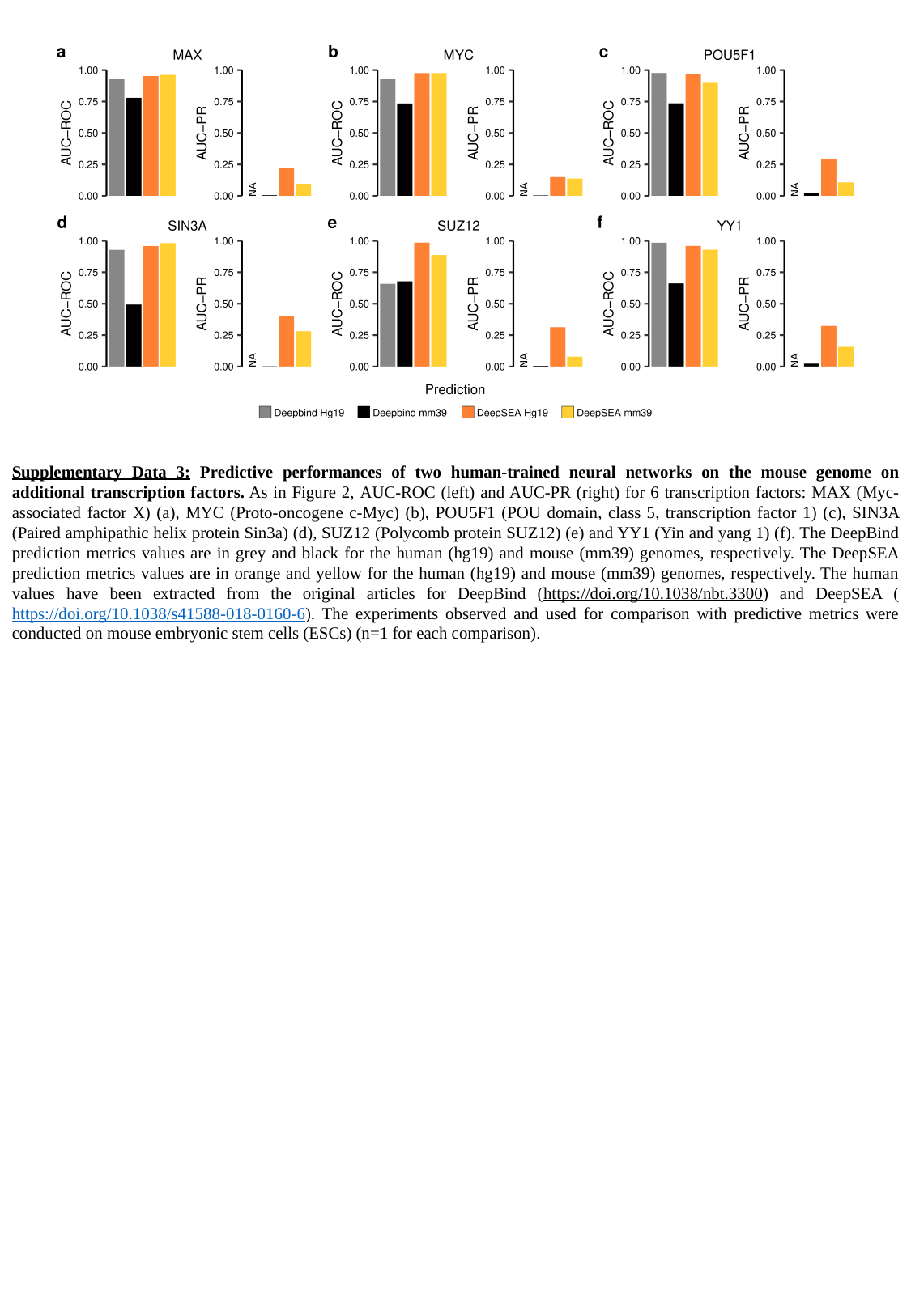

Supplementary Data 3: Predictive performances of two human-trained neural networks on the mouse genome on additional transcription factors. As in Figure 2, AUC-ROC (left) and AUC-PR (right) for 6 transcription factors: MAX (Myc-associated factor X) (a), MYC (Proto-oncogene c-Myc) (b), POU5F1 (POU domain, class 5, transcription factor 1) (c), SIN3A (Paired amphipathic helix protein Sin3a) (d), SUZ12 (Polycomb protein SUZ12) (e) and YY1 (Yin and yang 1) (f). The DeepBind prediction metrics values are in grey and black for the human (hg19) and mouse (mm39) genomes, respectively. The DeepSEA prediction metrics values are in orange and yellow for the human (hg19) and mouse (mm39) genomes, respectively. The human values have been extracted from the original articles for DeepBind (https://doi.org/10.1038/nbt.3300) and DeepSEA (https://doi.org/10.1038/s41588-018-0160-6). The experiments observed and used for comparison with predictive metrics were conducted on mouse embryonic stem cells (ESCs) (n=1 for each comparison).

### Slide 4
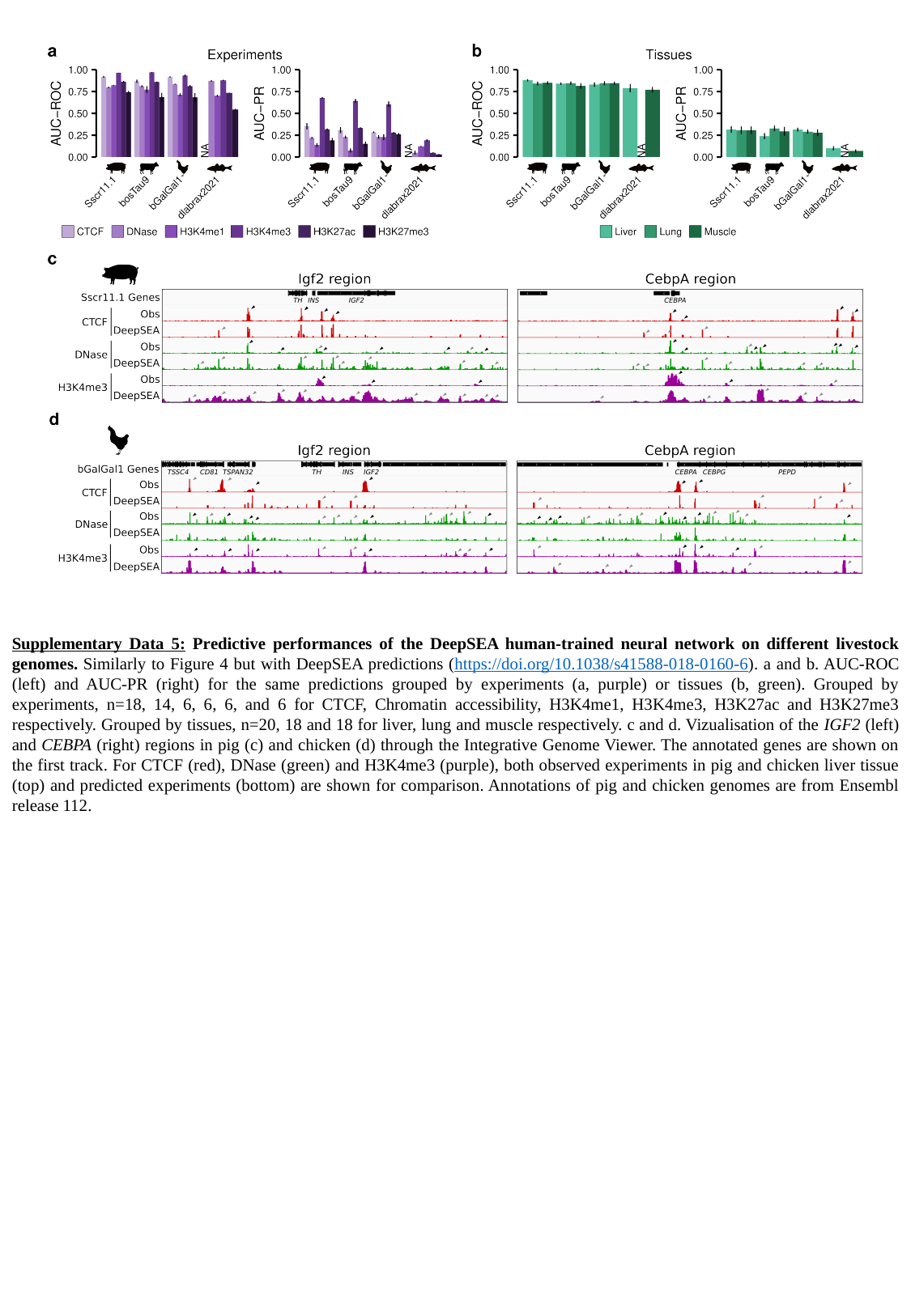

Supplementary Data 5: Predictive performances of the DeepSEA human-trained neural network on different livestock genomes. Similarly to Figure 4 but with DeepSEA predictions (https://doi.org/10.1038/s41588-018-0160-6). a and b. AUC-ROC (left) and AUC-PR (right) for the same predictions grouped by experiments (a, purple) or tissues (b, green). Grouped by experiments, n=18, 14, 6, 6, 6, and 6 for CTCF, Chromatin accessibility, H3K4me1, H3K4me3, H3K27ac and H3K27me3 respectively. Grouped by tissues, n=20, 18 and 18 for liver, lung and muscle respectively. c and d. Vizualisation of the IGF2 (left) and CEBPA (right) regions in pig (c) and chicken (d) through the Integrative Genome Viewer. The annotated genes are shown on the first track. For CTCF (red), DNase (green) and H3K4me3 (purple), both observed experiments in pig and chicken liver tissue (top) and predicted experiments (bottom) are shown for comparison. Annotations of pig and chicken genomes are from Ensembl release 112.
